# Eusocial reproduction selects for longevity

**DOI:** 10.1101/2025.03.25.645350

**Authors:** Rafael D’Andrea, Charles D. Kocher, Bonnie Skiena, Bruce Futcher

## Abstract

Animals such as bees, ants, wasps, termites, and naked mole-rats live in colonies in which a single queen is the only female reproductive, an arrangement known as eusociality. Eusocial animals are known for their remarkably long lifespans. It has been argued that longevity becomes selected when queens are shielded from “external mortality”. While such protection may contribute, we find a deeper reason: the eusocial reproduction strategy itself inherently creates selection for long lifespans. Lifespans typically reflect two processes: the baseline risk of death and the rate at which this risk increases with age. Each is a parameter in the Gompertz mortality equation. We show that the mathematical properties of eusocial reproduction lead to slowly-growing, older populations where selection acts more strongly on the rate at which risk increases than on the baseline risk. In addition, we show that channeling reproduction through a single female also selects for longevity, which we term the "queen effect". Thus, the dynamics of eusocial reproduction select for longer lifespan. More broadly, these results show that reproductive structure and population growth dynamics can fundamentally shape selection on lifespan, with implications outside eusocial systems as well.

## Introduction

Humans and other animals have an exponentially increasing risk of death with age. In humans, the risk of dying increases by about 8% per year after sexual maturity, such that the risk of dying doubles about every 9 years. This exponential increase in mortality with age was described by Gompertz [1], and is typical of many animals [2-6], although there are exceptions [7]. The mechanism behind exponentially-increasing mortality is unknown.

Among these apparent exceptions is the naked mole-rat, *Heterocephalus glaber*. This unusual mammal lives in underground burrows in Africa, eating tubers, and sometimes eaten by snakes [8]. Naked mole-rats reproduce eusocially: that is, a colony has only one reproductive female. Naked mole-rats have strikingly long lifespans of up to 40 years [9-11]. Mammalian life spans are generally well-correlated with body size [12, 13], but naked mole-rats are roughly mouse-sized mammals that live about five-times longer than expected [14, 15]. With a lifespan of about 40 years and a mass of about 35 grams, naked mole-rats live at least 10 times longer than *Mus musculus*, which has a lifespan of 1.5 to 4 years and a mass of about 20 grams. Recent studies suggest that naked mole-rats have a flat Gompertz curve with a slope near zero. That is, naked mole-rats apparently do not face a significantly increased risk of dying with older age [11, 16, 17].

Strikingly, other eusocial animals also have remarkably long lifespans [18-21]. The 2,600 known species of termites are eusocial, and all have very long lifespans. For example, queens of the species *Macrotermes michaelseni* have a mean lifespan of about 18 years [18]. The mean lifespan of queens across termites is 11.5 years, 10 years in ants, and 5.6 years in honeybees. By comparison, the mean lifespan of related solitary species is about 0.1 years [19]. Lifespans in queens of eusocial stinging wasps range from about 100 to 10,000 days, compared to 10 to 100 days among solitary stinging wasps [22]. Although workers live about one tenth as long as queens, they still commonly live 10-fold longer than their equivalents in non-eusocial species [22]. For instance, in a eusocial honeybee species where queens live about 1,080 days, workers live about 180 days. In contrast, sawflies, a non-eusocial insect from another suborder of *Hymenoptera*, have lifespans of 7 to 9 days. We are not aware of any exceptions to the rule that eusocial species have long lifespans, and this motivated us to consider whether eusocial reproduction, as such, might promote long lifespan.

Many researchers have studied this association between long lifespan and eusocial reproduction. It has been suggested that long lifespan causes eusocial reproduction [23, 24], and also that eusocial reproduction causes long lifespan [19]; (see [25] for a discussion of this large literature). The general argument for the latter is that species with eusocial reproduction have low extrinsic mortality (i.e., mortality from outside forces, such as predators, rather than internal factors such as cancer), which enables long lifespans and the selection of mutations promoting longevity. Specifically, it has been suggested that eusocial species are often found in a protected environment—a hive, or nest, or burrow—where individuals, especially the queens, are protected from extrinsic dangers [19, 26-28]. Oster and Wilson described a colony of eusocial insects as “a factory within a fortress” [29]. However, it has been questioned whether low extrinsic mortality is a complete explanation [30], and the interplay between extrinsic mortality, density-dependent mortality, and other forces may be relevant [31-33].

Here we provide a different explanation. We show that the dynamics of eusocial reproduction causes selection for long lifespan regardless of sheltering from harsh environments. Eusocial reproduction leads to populations with higher average ages, and also to slower population growth, which reduces early mortality from resource limitation. These select for longevity. In addition, we suggest that funneling reproduction through a single female, the queen, may on its own select for longevity.

### Rationale. Why might eusocial reproduction favor longevity?

Figure 1 compares population growth under two reproductive strategies. Temporarily ignoring death and resource constraints, the number of offspring grows exponentially with time under regular reproduction, but linearly under eusocial reproduction. The median age in this regular-reproduction population is 0 (the age of newborns), while in the eusocial population, the median age is (queen lifespan/2). Thus, before considering death and resource limitation, eusocial populations will tend to be smaller and skew older. As a shorthand, we will refer to eusocial reproduction as linear reproduction, and regular reproduction as exponential reproduction.

**Figure 1.**
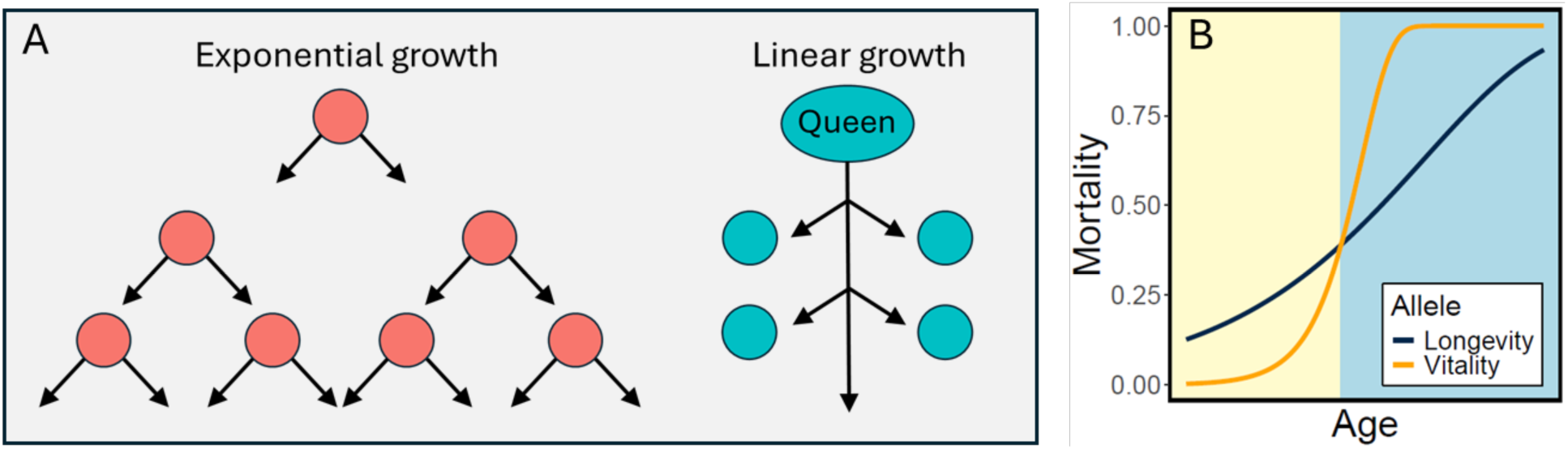
Comparative Dynamics of Exponential and Linear Reproduction. **A.** Diagram of population growth in normal (exponential) vs eusocial (linear) reproduction. When all individuals reproduce, the population grows exponentially, and median ages are young. When only the queen reproduces, the population grows linearly, and median ages are older. **B.** The Gompertz mortality curve for hazard *h* at age *n* is *h*(*n*) = *αe*^*βn*^ + *γ*. Age-dependent mortality is shown for animals differing in two genetic alleles specifying different values for the Gompertz parameters α and β. Lower values of α (high “Vitality”) benefit younger individuals (left side, yellow), while lower values of β (high “Longevity”) benefit older individuals (right side, blue). Parameters: Longevity allele (*α* = 0.125, *β* = 0.15), Vitality allele (*α* = 0.002, *β* = 0.6).

Age-dependent mortality in humans and other animals can be described by the Gompertz-Makeham law, where the hazard function at age *n* is

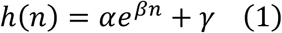

Parameter *γ* reflects the age-independent component of mortality. Parameter *α* sets mortality at age 0; that is, the baseline death rate before senescence, reflecting a young individual’s strength, health, and durability—a trait we call “Vitality”. The *β* parameter determines the steepness of the mortality curve, or how quickly mortality increases with age; lower values of this parameter increase “Longevity”. Essentially, an animal’s vulnerability to death increases exponentially with time at a rate set by longevity. For instance, survival might decrease with a half-life of 9 years, as in humans. Increased longevity would increase the half-life of decline (e.g., to 10 years) and so lead to a shallower decrease. Our central argument is that depending on an animal’s life history, selection may act more strongly on vitality (i.e., overall durability), or on longevity (i.e., the half-life of decline), and that for eusocial animals, selection acts strongly on longevity.

In the Gompertz curve (Fig. 1B), high α or β lead to high mortality (more accurately, high hazard). For convenience, we henceforth focus on the inverse, survival. In what follows, our “Vitality” parameter is related to the inverse of α, and our “Longevity” parameter is related to the inverse of β. That is, high Vitality (low α) and high Longevity (low β) can both lead to high survival.

Fig. 1 illustrates two intuitive reasons why linear reproduction selects for longevity:

1. **Age distribution.** The median age in an exponential population will be younger than in an otherwise equivalent linear population. Therefore, an exponential population tends to live on the left side of Fig. 1B, where vitality is preferred, while a linear population tends to live on the right side, where longevity is preferred. Thus, selection for longevity will have a greater effect on a linear population.
2. **Resource pressure.** Growing populations will reach an equilibrium size at the carrying capacity of their environment. Exponential populations, with their high rate of reproduction, will push hard against this population cap, and resource limitation will be a major source of death, often at young ages, due to lack of resources rather than to senescence. In contrast, linear populations, with a smaller capacity for reproduction, will push less hard against the population cap, and senescence will be relatively more important. The result will be that exponential populations will have shorter lifespans and select for vitality, while linear populations will have longer lifespans and select for longevity.
3. **Queen effects**. In addition, and not shown in Fig. 1, less intuitive “queen effects” arise from the channeling of reproduction through a single female. For example, in a linear population, a mutant allele for increased longevity of the queen will spread because it will produce significantly more offspring than its wildtype allele. In an exponential population, this allele would have a smaller advantage, since offspring of older females would form only a small portion of the next generation due to the much greater number of younger reproductive females. Other potential “queen effects” are described in the Discussion.

### Approach. Study the relative selection for vitality or longevity

We compare the fates of two competing alleles of a gene in a population. Each allele specifies values for both vitality (V) and longevity (L) (i.e., for the inverses of the α and β parameters of the Gompertz curve). The “Vitality” allele has a relatively higher value for vitality at the expense of lower longevity, while the “Longevity” allele has the opposite properties. This was inspired by G.C. Williams idea of “antagonistic pleiotropy” [34]. Given a pair of alleles with different but nearly balanced vitality and longevity values, we compete that pair of alleles under exponential reproduction, and under linear (i.e., eusocial) reproduction. We show that for a single pair of nearly balanced alleles, the vitality allele consistently “wins” under exponential reproduction, while the longevity allele consistently “wins” under linear reproduction, i.e., that longevity is favored under linear reproduction.

Although reproduction within a colony is linear, colonies themselves reproduce exponentially. We show in Results 2 (Colony Reproduction) that this colony-level exponential growth does not override the effects of linear reproduction at the individual level. This is because exponential colony reproduction selects on colony-level traits (e.g., seeding and collapse), which are only indirectly related to phenotypes at the individual level.

## Results

Results are in three parts. First, we describe selective forces in the context of reproduction of individuals. We address this with an analytical approach (Results 1a) and with a numerical approach (Results 1b). Next, we describe the reproduction of colonies and how this affects selective forces on individuals. Again, we address this with an analytical approach (Results 2a), and a numerical approach (Results 2b). Finally, we describe a “Queen effect” promoting longevity uniquely in eusocial animals (Results 3).

### Results 1. Individual Reproduction

#### 1a. Analytical Model

We mathematically modeled natural selection acting on populations with linear or exponential reproduction to explore the relative fitness advantages of longevity in each scenario. In brief, we developed an age-structured discrete-time model of these types of populations to determine fixation in a competition between two alleles. We derived a relative fitness function for these competitions, Δ*F*, whose sign determines the winning allele:

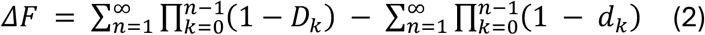

Here, *D*_*k*_ and *d*_*k*_ are the probabilities that individuals of age *k* will die before they reach age *k* + 1 for each allele, computed from the Gompertz hazard function. For mathematical details, see Methods and the SI.

Many different combinations of vitality and longevity in an allele can yield the same fitness, since gains in one can offset losses in the other. We analytically mapped the combinations of vitality and longevity that maintain constant fitness to generate fitness nullclines using Eq (2). As Fig. 2 shows, the calculated nullclines for exponential and linear populations have very different slopes. In the exponential population, a small change in vitality requires a large change in longevity to offset it, whereas in the linear population, the same change in vitality can be offset by a smaller change in longevity—in other words, longevity is relatively more potent in the linear population. In comparative terms, this suggests that exponential populations “care” more about vitality while linear populations “care” more about longevity.

**Figure 2:**
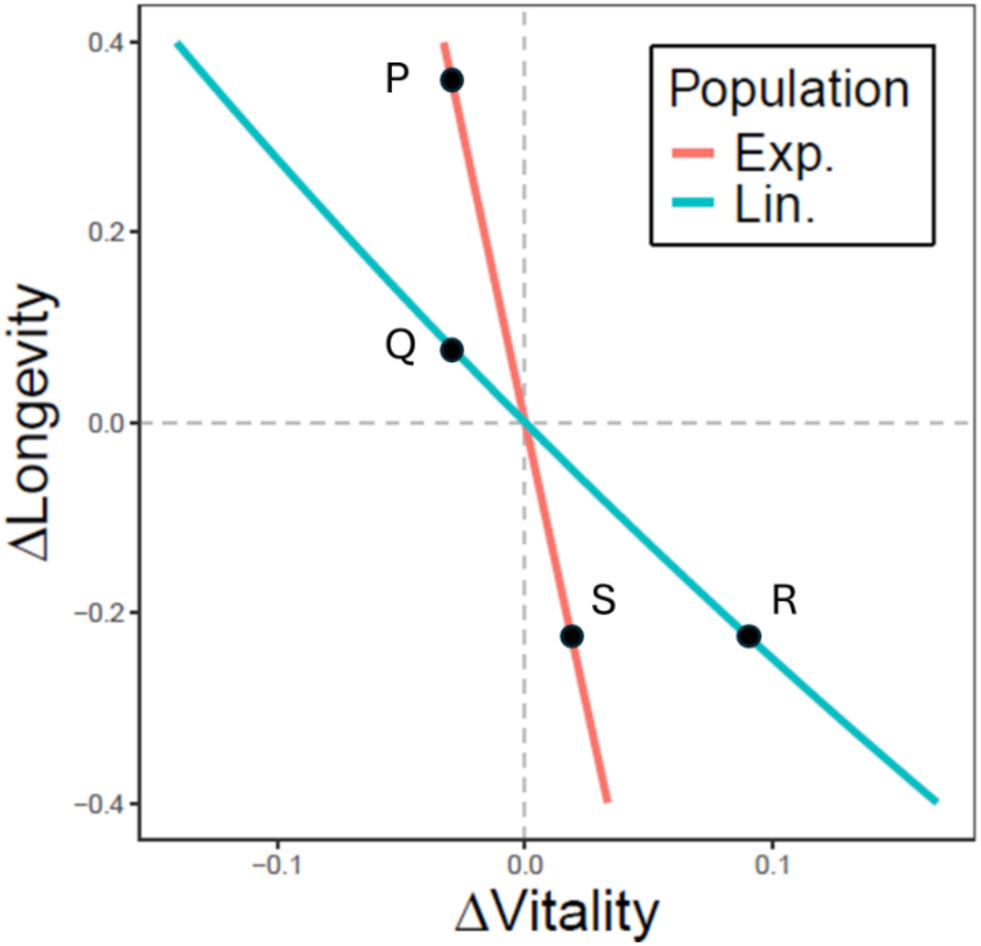
Fitness nullclines (Δ*F* = 0) for alleles balancing vitality and longevity in exponential (red) and linear (blue) populations. Slopes indicate the relative strength of selection on each trait: steeper slopes in the exponential population require larger gains in longevity to offset losses in vitality relative to the origin (compare P vs. Q), indicating higher sensitivity to vitality. In contrast, shallower slopes in the linear population require larger gains in vitality to offset losses in longevity (R vs. S), indicating higher sensitivity to longevity.

#### 1b. Numerical Model

The analytical treatment requires simplifying assumptions, bypasses the genetics of reproduction, and gives no opportunity to examine life-history parameters such as time to sexual maturity, probability of reproduction, litter size, the intensity of resource limitation, and patterns of genetic dominance. Therefore, we developed an agent-based model, the “NMR Model” to simulate dynamics in populations with exponential or linear reproduction.

Using the NMR Model, we competed a vitality allele against a longevity allele—with the former providing 10% higher vitality but 10% lower longevity compared to the latter, with each initially comprising 50% of the population. Specifically, we used a vitality allele with the parameters (V=1000, L=9) and a longevity allele with parameters (V=900, L=10) (Methods). This longevity allele fixes (i.e. reaches 100% frequency) more often in the linear population. Across 1000 simulations of each population type, the longevity allele fixed in 58% of the linear populations but only 6% of the exponential populations (Fig. 3, Supplemental Table 1).

**Fig. 3.**
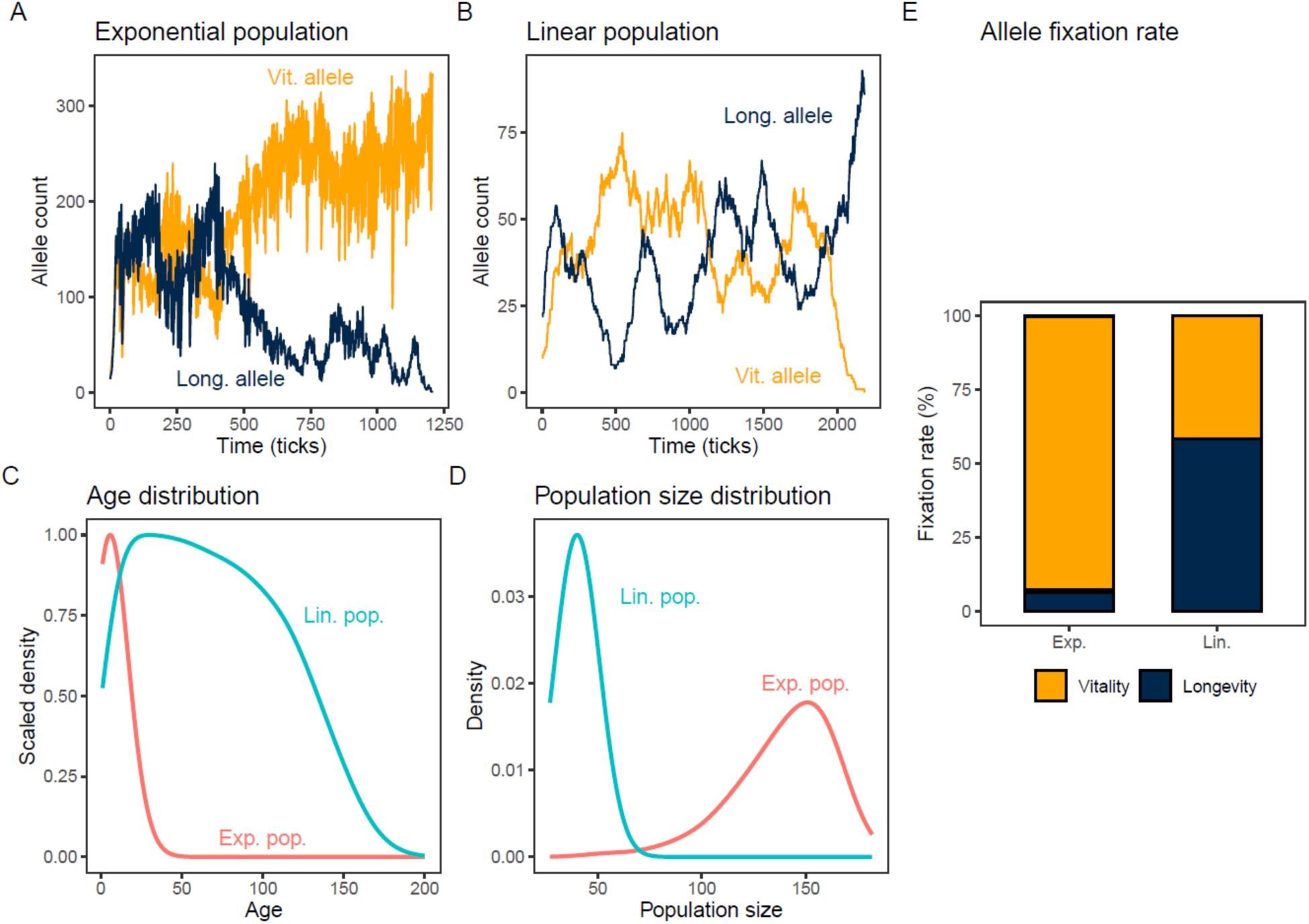
The Longevity allele is preferred in linearly-reproducing populations. A dominant Vitality allele (V =1000, L=9, yellow) competed against a recessive Longevity allele (V=900, L=10, dark blue) in the NMR numerical model. Similar results were obtained when dominance was reversed. **A**. Time series of the frequencies of the longevity and vitality alleles for an example simulation of an exponential population. **B**. Time series of the frequencies of the longevity and vitality alleles for an example simulation of a linear population. **C** and **D**. Age distributions and population size distributions in the exponential (red) and linear (blue) populations, over 1000 simulations. **E**. Relative fixation rates of competing vitality (yellow) and longevity (dark blue) alleles over 1000 simulations each of exponential and linear populations.

This relative advantage of the longevity allele in the linearly growing population held across a wide range of scenarios of the other parameters (dominance, time to sexual maturity, litter size, resource limitation, etc.). Notably, when all parameters other than vitality and longevity were identical between the two reproductive strategies, we found no scenario in which their preferences reversed—exponential reproduction always preferred vitality, linear reproduction always preferred longevity. Exponential populations were consistently much larger and composed of much younger individuals than linear populations (Fig. 3C, 3D). This is primarily because rapid population growth in the exponential case quickly leads to resource limitation, bringing death upon individuals of all ages. This pattern holds across a range of parameter values and multiple resource limitation scenarios.

We note that the difference in relative success of the two alleles between the two population types is likely even larger than suggested by these results. The linear population has a very small effective population size (*N*_*e*_ ≲ 10) due to the genetic bottleneck caused by a single reproductive female. This causes powerful genetic drift in the linear population, such that even disfavored alleles can fix frequently. The difference in effective population sizes between the two kinds of populations makes it difficult to calculate comparable selection coefficients. However, the sign of the selection coefficient towards longevity in the linear population is clear (Fig. 3E).

### Results 2. Colony Reproduction

Here we show that the super-imposition of the exponential reproduction of colonies on the linear reproduction of individuals does not affect our conclusions—linear (eusocial) reproduction still selects for longevity.

#### 2a. Analytical Model: Colony Reproduction

The results above show that individuals reproducing linearly experience a stronger selective force for longevity relative to individuals who reproduce exponentially. However, colonies reproduce exponentially, since every colony is capable of spawning other colonies. Arguably, this higher-level exponential reproduction might weaken or nullify the selection towards longevity observed above. However, that is not the case.

As discussed in detail in Methods, the selection for longevity identified above acts on the age dependence of death rates of individuals. When the same logic is applied to exponentially-reproducing colonies, it is the age dependence of the colony’s death (collapse) rate that experiences selection, rather than the death rates of the individuals within the colony. In other words, selection acting at the colony level operates on colony-level variables, which are only indirectly affected by the phenotypes of individual colony members. Thus, exponential reproduction of colonies does not imply any particular selective force on the longevity of individuals and therefore does not negate the selective effect of linear reproduction acting on individuals.

The question remains whether colony reproduction influences the selective force in other ways. In general, between-group selection can be different from within-group selection [35-37]. In Methods, we prove for a simplified scenario of colony reproduction that the relative fitness function for the colony competition is the same as Eq (2). This demonstrates that at least in simple cases, colony reproduction does not impact our conclusions.

Despite the simplifications, this finding suggests that the preference for longevity by linearly growing populations might be preserved more generally, colony reproduction notwithstanding. For example, Wilson [37] discussed two limiting cases: pure individual selection (analogous to our individual reproduction), and pure group selection, where each group consists entirely of a single allele type (analogous to our colony reproduction). As the strength of group selection increases, the system transitions from the pure individual case toward the pure group case (see Fig. 1 of that paper). All outcomes fall between these two extremes. Our analytical results show that selection in both of our limiting cases is identical; therefore, for any intermediate strength of group selection, the result should match that in the individual case. Although Wilson’s model differs from ours in several details, we expect the underlying principles to be similar.

#### 2b. Numerical model of Exponential Colony Reproduction

To ask whether reproduction of colonies affected the selection for individual longevity, we extended the numerical model of Results 1b to encompass exponential reproduction of colonies. Within a colony, population growth follows linear reproduction as in 1b above. Colonies can seed new colonies exponentially, and existing colonies can die. Analogously to 1b above, a logistic limit was placed on colony reproduction, such that when the total number of colonies becomes large, it becomes exponentially more difficult to form new colonies. Here, there is no analogue for exponentially reproducing populations, for which colonies are undefined.

We wrote two versions of the model to mimic features of two known eusocial lifestyles. The first was “Ant mode”, which mimics some features of ant colony reproduction [38]. In Ant mode, new colonies are founded by a queen who has an infinite supply of semen from a “mating flight”. Upon founding a colony, the queen produces offspring indefinitely, using stored semen. When the colony reaches a “Threshold Colony Size”, it sends out “mating flights” of new reproductives to found new colonies. When the queen dies, the entire colony dies.

The second was “Mole-rat mode”, which mimics some features of naked mole rats [9, 39-43]. Queens mate within colonies. When a queen dies, she is replaced by another female from the same colony. When the colony reaches “Threshold Colony Size”, it sends out pioneers to found new colonies. In Mole-rat mode, each colony has a low probability (typically 0.045 per time interval) of encountering a “Disaster”, which has a high probability of destroying the entire colony.

We tested whether reproduction of colonies reverses the preference of eusocial reproduction for longevity. We again competed a vitality allele (V=1000, L=9) against a longevity allele (V=900, L=10), but with colony reproduction superimposed on individual linear reproduction (Fig. 4). As expected from the analytical argument, the longevity allele continued to win over the vitality allele (Fig. 4C), both in Ant and Mole-rat Mode. In fact, the preference for the longevity allele appears stronger with colony reproduction (Fig. 4C). This is likely due to the larger effective population size relative to the individual-reproduction model--the advantage of the longevity allele is no longer dampened by genetic drift.

**Figure 4.**
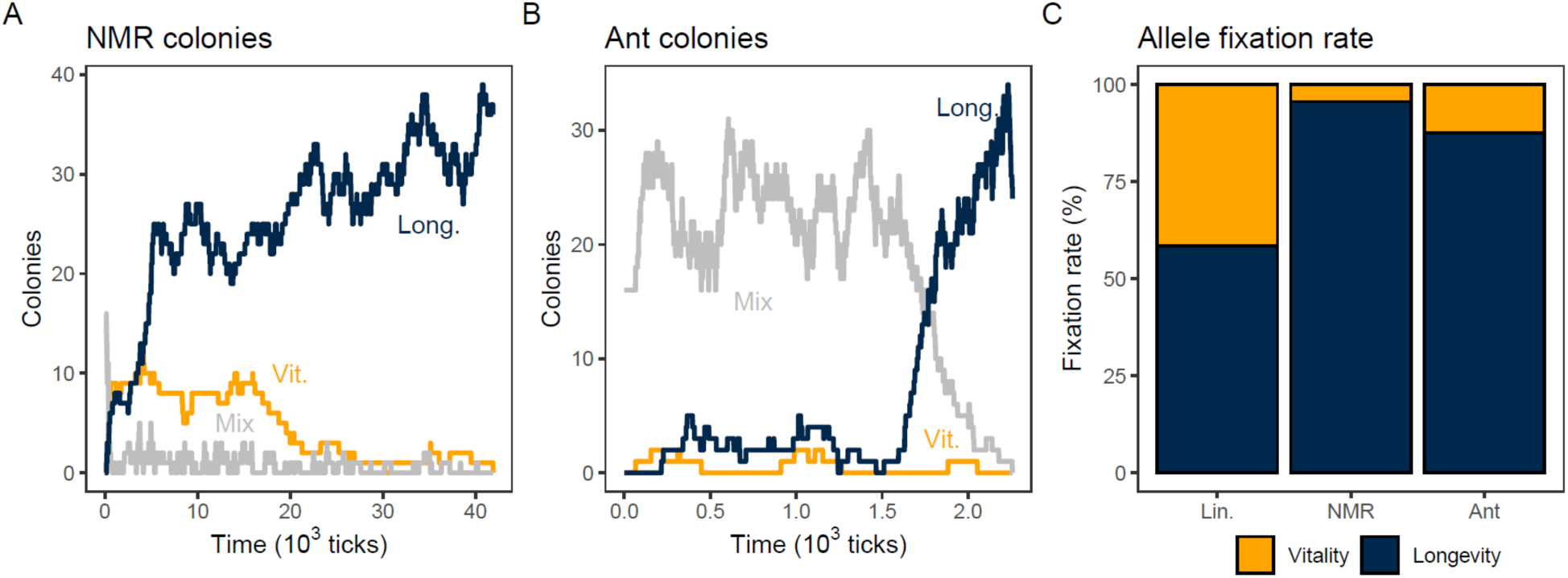
The longevity allele is preferred even when colonies reproduce. Two modes of colony reproduction (Mole-rat mode, “NMR”, and Ant mode, “Ant”) were super-imposed on linear individual reproduction in the NMR numerical model. A dominant Vitality allele (V =1000, L=9, yellow) was competed against a recessive Longevity allele (V=900, L=10, dark blue). **A**. Time series of the frequencies of the longevity and vitality alleles for an example simulation of a Mole-rat colony population. **B**. Time series of the frequencies of the longevity and vitality alleles for an example simulation of an Ant mode population. **C**. Relative fixation rates of competing vitality (yellow) and longevity (dark blue) alleles after 1000 simulations each of Mole-rat and Ant mode, compared to the fixation rate of the same alleles in individual reproduction (“Lin.”, same data as Fig. 3E).

The colony reproduction models had fourteen adjustable parameters in addition to those of the individual-reproduction model. We explored all of these individually, and several in combination. While these parameters slightly affected the degree of preference for longevity, none reversed it.

To further explore differences in selection between the individual and colony reproduction models, we tested two new longevity alleles with weakened longevity. The (V=900, L=9.2) longevity allele loses to the (V=1000, L=9) vitality allele under exponential reproduction, but ties with it under linear reproduction; that is, (V=900, L=9.2) is a neutral allele under linear reproduction. The (V=600, L=9.3) longevity allele is even weaker and loses to the vitality allele even under linear reproduction. Results for these weakened longevity alleles in colony reproduction are shown in Fig. 5. In both Ant and Mole Rat Mode, the success of the weakened alleles in colony reproduction was directionally comparable to their success in individual reproduction, again arguing that colony reproduction does not alter the selection for longevity.

**Figure 5.**
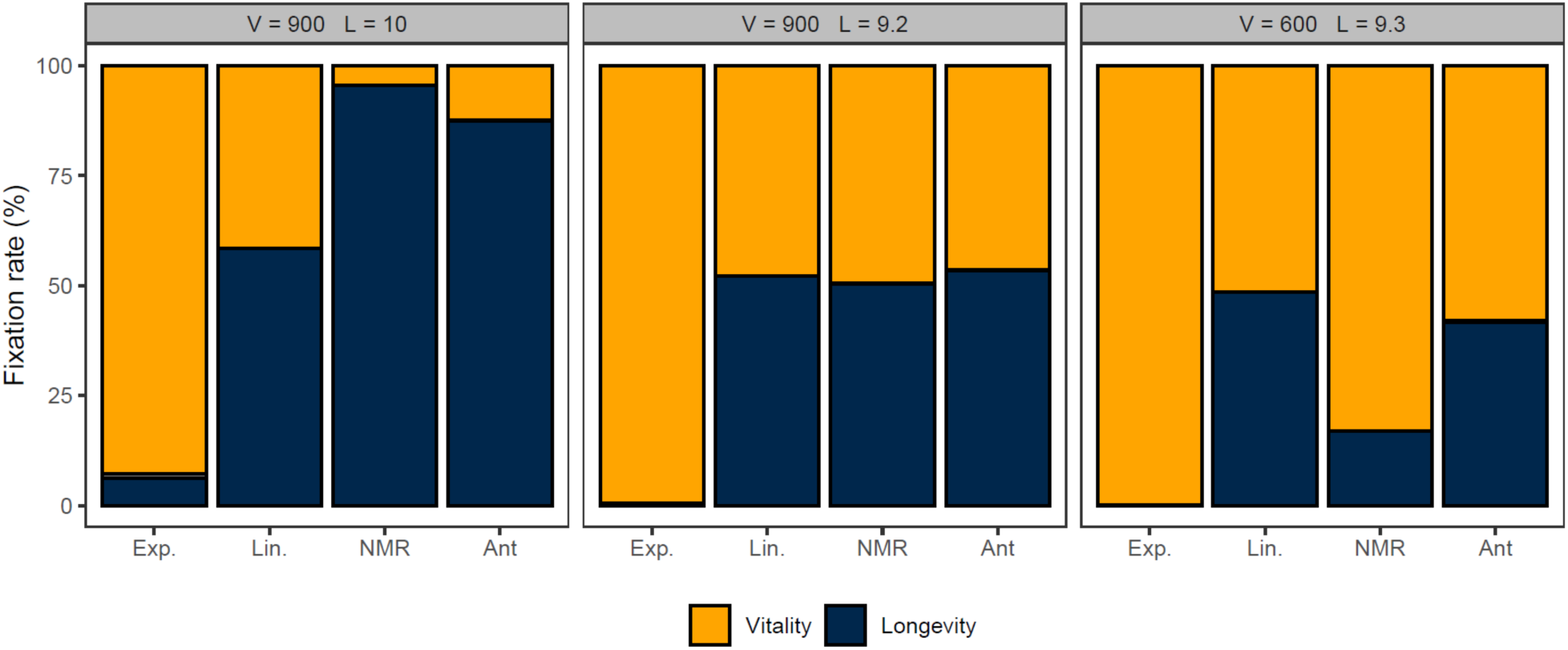
Comparison of strong, neutral, and weak longevity alleles. The standard vitality allele (V=1000, L=9) was competed with strong (V=900, L=10), neutral (V=900, L=9.2) and weak (V=600, L=9.3) longevity alleles, in individual reproduction (exponential, “Exp.”, and linear, “Lin.”) and in both modes of colony reproduction (Mole-rat mode, “NMR”, and Ant mode, “Ant”) super-imposed on linear reproduction, using at least 1000 runs each of the NMR model. There are rare cases where neither allele fixed (thin grey segments, usually too thin to be visible, but see left-most bar). Data for the strong allele is the same data as in Fig. 4C and Fig. 3E, and is re-shown for comparison. Exact results are shown numerically in Table S1 of Supplementary Information, Appendix 3.

The weakest longevity allele (V=600, L=9.3) does worse under colony reproduction than under linear reproduction (Fig. 5), again suggesting that the magnitude of results under individual linear reproduction is dampened by the small effective population size. Indeed, we did not observe a shift in the neutral allele for Mole-rat mode compared to the individual Linear population, as the fixation of each on the (V=900, L=9.2) allele was consistent with neutrality (using 95% Wald confidence intervals; see the SI). For Ant mode, this allele was not neutral (p_fix_ = 0.534±0.031); longevity preferentially fixed, suggesting that the Ant mode of colony reproduction slightly increased the preference for longevity.

Combined, these results show that colony reproduction has a large impact on the effective population size but only a small impact on the selection coefficient between the vitality and longevity genes—if anything, it increases the preference for longevity.

#### Results 3. Queen Effects

We can adjust parameters so that a linear population matches the size and age structure of an exponential population. For example, increasing the queen’s fecundity in the linear population boosts population size (and thereby shortens lifespans via resource limitation), whereas delaying sexual maturity, reducing the probability of reproduction, or shrinking litter size in the exponential population has the opposite effect. Indeed, natural variation in these life history traits across eusocial and non-eusocial populations can modulate the selection for longevity. However, eusocial and non-eusocial populations still differ in that the linear population channels reproduction through a single female.

To isolate the effects of channeling reproduction through a queen, we created a second numerical model (the “Queen effects-only” model), and adjusted parameters (see Materials and Methods) to equalize population size and age distribution between linear and exponential populations. This allowed us to test how removing these differences influenced the competition dynamics of the Vitality and Longevity alleles. Here, a wildtype allele is invaded by a mutant, initially at frequency *p*_0_, that conferred higher vitality but lower longevity. We measured the mutant’s fixation rate *f* and calculated its fixation bias FB = *f*/*p*_0_ − 1, where FB = 0 indicates a neutral mutant whose vitality advantage exactly offsets its longevity disadvantage (i.e., it is on one of the nullclines of Fig. 2). To identify this neutral mutant in each population type, we varied the vitality advantage while keeping longevity loss fixed and determined the threshold at which FB changes sign. Our results show that exponential populations require less of a vitality advantage to balance the longevity cost compared to linear populations (Fig. 6). Thus, even in the absence of age and resource pressure effects, the linear population shows a persistent, albeit smaller, predilection for longevity relative to the exponential population.

**Figure 6:**
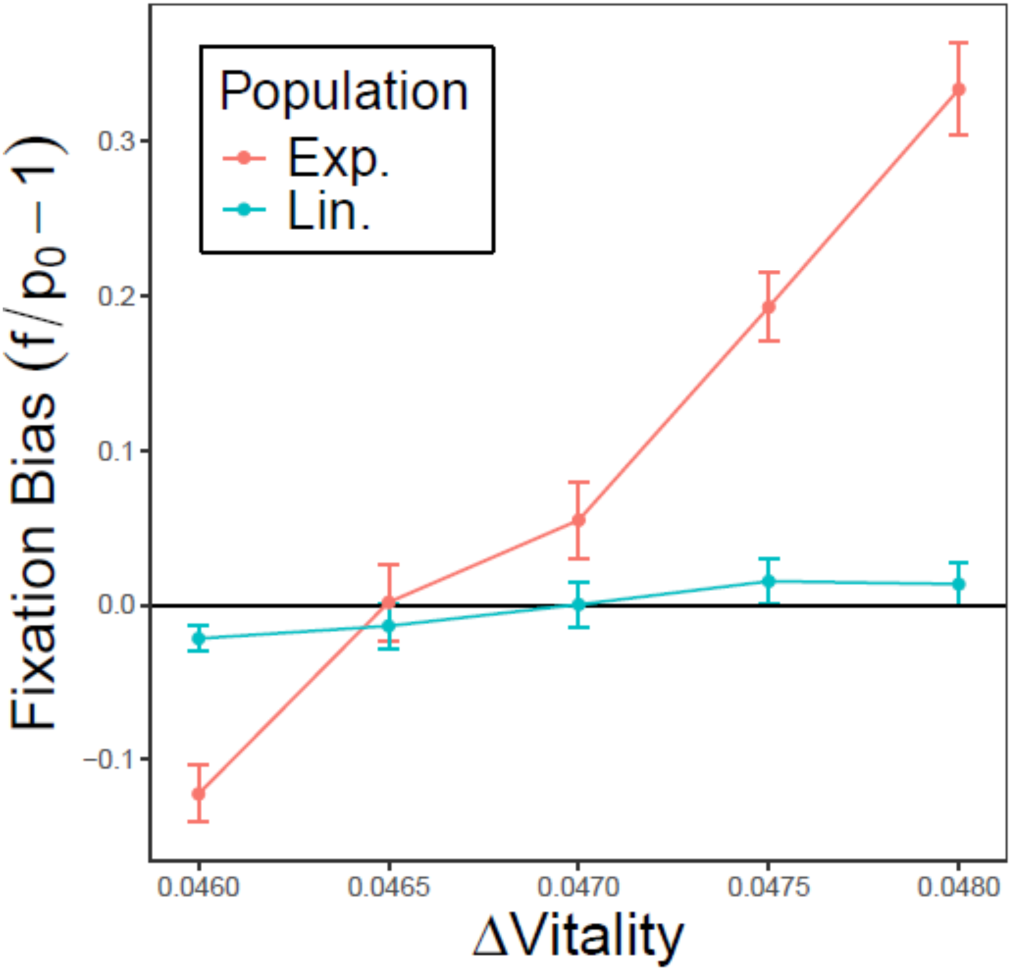
Fixation bias of an invading allele with higher vitality but lower longevity in exponential (red) and linear populations (blue). The x-axis shows the vitality difference between the invader and wildtype, while the y-axis shows f/p0 - 1, where f is the observed fixation probability and p0 is the initial frequency of the invader. The intersection of each curve with y = 0 marks the vitality advantage required for the lower-longevity invading allele to match the wildtype. While both selection and effective population size influence the steepness of the curves, only selection affects the intersection point. Results from the “Queen effects-only” numerical model.

Note that while the intercepts of the curves with the neutral line indicate differences in selection for longevity between the two population types, their slopes are influenced by effective population size. The smaller effective size of the linear population results in a shallow slope due to drift, effectively dampening the apparent strength of selection.

## Discussion

Our findings suggest that, all else being equal, a switch in reproductive strategy from regular (exponential) to eusocial (linear) would change population structure: the population size would decrease, causing less pressure against the environmental resource cap, and the average age would increase. These changes would cause selection for increased longevity, especially in the queen.

These selective forces are not limited to eusocial reproduction. Life history strategies that slow down exponential reproduction tend to select for longevity [44]. That is, the effects of linear reproduction on population size and age distribution can be mimicked by exponentially-reproducing species. For example, K-selected traits such as increased age at sexual maturity, smaller litter size, and longer interbirth intervals would all cause smaller, slower-growing populations and lower resource pressure, higher average ages, and hence increased selection for longevity [44], by the same forces we have described above. Linear (eusocial) reproduction is arguably the limiting case of slow exponential reproduction, where both lead to older populations, reduced resource pressure, and stronger selection for long lifespan (see Vries et al. [32] for a similar argument). An extreme example of slow exponential reproduction is the Greenland shark (*Somniosus microcephalus*): it reaches sexual maturity at around 150 years, has an 18-year pregnancy, and is the longest-lived vertebrate known, with a lifespan of about 400 years [45, 46].

Reproductive skew towards females in exponential populations could also mimic eusociality for these purposes [47]. When a small proportion of the females produce a large proportion of the children, this is partway to eusocial reproduction. Indeed, many animals with female reproductive skew have been reported to have notably long lifespans. Examples could include the spotted hyena (*Crocuta Crocuta*), the meerkat (*Suricata suricatta*), the common marmoset (*Callithrix jacchus*), the yellow-bellied marmot (*Marmota flaviventris*) and the common eider duck (*Somateria mollissima*). However, there is substantial variation in the literature values for the lifespans of these animals and (for comparison) for their close relatives with less reproductive skew. By contrast, a skew in male reproduction does not produce the same effects, since males generally do not face the same limitations in offspring production as females—expensive gametes, egg laying and incubation, pregnancy, maternal care, etc. In fact, male reproductive skew is often associated with short life span because of the costs of competing with other males for mates.

Beyond age- and size-related effects, the channeling of reproduction through a single individual can generate “queen effects” that are unique to eusociality. Queen effects can result from at least three different mechanisms. First, an invading gene that increases queen longevity has a fitness advantage over the wildtype, as the queen monopolizes reproduction. In an exponentially reproducing population the same gene would confer a smaller advantage because the long-lived female’s offspring constitute a small fraction of the total population. Second, under linear reproduction, the queen’s alleles tend to fix due to the small effective population size. If the queen outlives other females, fixation becomes even more likely, and the gene may spread even without increasing population-level fitness. The effect is particularly pronounced when only a few males are reproductive, further decreasing N_e_, as seen in many eusocial species. By becoming a long-lived queen, an individual can effectively exclude competing alleles, ensuring the dominance of her own genetic lineage.

A third potential mechanism for a queen effect involves investment of resources in physical size and strength of the queen, improving her overall survival and reproductive capacity. In an exponential population, all females would have to be so endowed, incurring a larger resource cost. Once queen vitality is improved, the selective advantage of improved longevity increases. Queens that are larger, stronger, and longer-lived than worker females are well-documented in bees, ants, wasps, termites, and naked mole rats (e.g., [40, 48, 49]). The concept is vividly displayed in fiction [50]. Critically, whereas improved size and strength require resource investment and are thus limited to the queen, improved longevity might not require additional resources and thus might not incur a cost if expressed in the workers. Thus, longevity-promoting alleles that are initially selected in queens could come to be expressed throughout the population. In line with this reasoning, previous simulation work has shown that lifespan differences can evolve between queens and workers [51]. In fact, in the eusocial termites and in Hymenoptera and in naked mole rats, both workers and queens tend to outlive their non-eusocial kin, with queens exhibiting particularly extended lifespans (e.g. [22, 52, 53]).

There is substantial literature describing possible mechanisms for the long life of naked mole-rats. These mechanisms include increased hyaluronan [54], an activated INK4a-RB senolytic death pathway [55], elevated glycogen, elevated expression of HIF-1α, and reduced succinate levels [56], resistance to oxidative stress [57], an improved immune system [58], an improved apolipoprotein A-I [59], extreme tolerance to hypoxia [60], improved regulation of splicing [10], mitochondrial anti-oxidant defenses [61], more efficient DNA repair [62-64], improved proteostasis [65], better telomere maintenance [66, 67], and many others. Our work does not address mechanism, but provides an evolutionary explanation for the unusual longevity of the naked mole-rat and other eusocial species. From this perspective, the particular mechanisms are perhaps less important than the selective forces behind them. That there are so many putative mechanisms may suggest that when selection for longevity emerges, organisms avail themselves of many mechanistic responses, which argues against a single major cause of ageing.

Our perspective is consistent with life history theory [44, 68, 69] and can be framed as a life history argument. However, our approach is simple, relying on only a few population properties while producing large effects. Notably, we show that different life histories select differentially on the α (vitality) and β (longevity) components of the Gompertz-Makeham hazard function. For exponentially reproducing populations, our conclusions broadly match traditional life history predictions (e.g., “live fast, die young” vs. “slow and steady” strategies). However, the comparison with linear reproduction puts these strategies in a new light. Eusocial species such as ants, bees, wasps, and naked mole-rats do not fit neatly on the traditional “live fast” vs “slow and steady” axis, but when analyzed as here, appear molded by some of the same underlying forces as “slow and steady” species, albeit in a nonstandard way.

Our conclusions also align with previous research on the long lifespans of eusocial animals, which often attributes their longevity to lower extrinsic mortality, possibly due to sheltered environments such as hives or nests [19, 20]. While we also invoke a kind of reduced extrinsic mortality, we propose that it arises primarily from slower population growth, which eases resource competition, rather than from fortress homes.

We note that workers may increase queen fecundity by assisting her with offspring care, which our analyses did not consider. This effect likely saturates quickly with increasing colony size due to inherent limits on queen fecundity. However, such “worker effects” remain an interesting subject for future research.

In summary, we argue that eusocial reproduction leads to older populations and reduced resource pressure, which select for longevity. These same forces also operate on non-eusocial species and manifest when reproduction slows down via increased age at maturity and other life history traits. Uniquely, eusocial populations have “queen effects” that also select for longevity. That selection for longevity is evidently so successful over such a wide variety of organisms suggests that there are likely multiple mechanisms of senescence, and accordingly multiple mechanisms of evading it, which could differ across the tree of life. Lifespan would appear to be an evolvable trait, highly responsive to selection.

## Methods

### Analytical Model Individual Reproduction

We consider a contest between a reference allele A and a competing allele B within a single population or single colony. Alleles A and B differ in their relative values of vitality and longevity, which set parameters of the Gompertz equation. We track the number of individuals of each phenotype at each age following the discrete-time update equations:

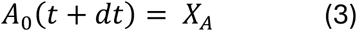

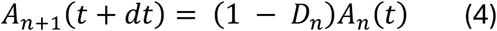

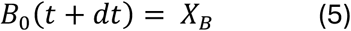

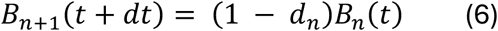

Here, *A*_*n*_(*t*) and *B*_*n*_(*t*) are the abundances of individuals of age *n* carrying each allele at time *t*, and *X*_*A*,*B*_ is the number of births of either type during a time interval *dt*, which depends on the abundances of both alleles at time *t* and the reproductive mode of the population (exponential or linear; we will give explicit forms later after developing some general results first). Individuals of age *n* survive to age *n* + 1 (measured in timesteps of *dt*) with probability 1 − *D*_*n*_for wild-type allele *A* and 1 − *d*_*n*_for allele *B*. The death probabilities *D*_*n*_ and *d*_*n*_, assumed density-independent for simplicity, increase with age according to the Gompertz-Makeham law for the age-dependent hazard rate *h*(*n*):

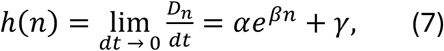

where the positive constants *α* and *β* determine the mortality of newborns and how quickly mortality increases with age, respectively. The concepts of vitality and longevity as used in this study are thus inversely related to *α* and *β*, respectively. The age-independent component *γ* is related to the carrying capacity of the population (see Appendix 1 in the Supplementary Information). Correspondingly, the death probability, or mortality, is

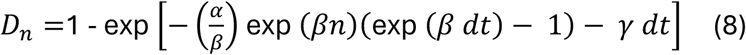

(see Equation S.18). The age-dependent component (*γ* = 0) of this quantity is plotted in Figure 1B, with *dt* = 1 scaling the time units.

We characterize our alleles based on different choices for *α* and *β*, and the tradeoff between longevity and vitality is implemented via a tradeoff between those two parameters. That is, we set

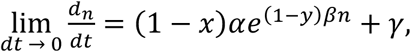

where *x* and *y*--the fraction change in *α* and *β*, respectively, from the reference wild-type--must be of opposite signs. If *x* > 0 and *y* < 0, then allele B confers higher vitality and lower longevity than allele *A*; when the signs are switched, the opposite is true.

The winner of the competition between alleles *A* and *B* is determined by the difference in their respective finesses. The result of a competition between the two alleles is given by the steady-state populations of A and B as dictated by the discrete-time update equations (3)-(6); there are only four options: (i) Both A and B have zero population at steady-state (trivial solution, which we will not address), (ii) both A and B have non-zero populations at steady-state (co-existence steady-state), (iii) only A is non-zero (competitive exclusion, with A more fit), and (iv) only B is non-zero (competitive exclusion, with B more fit). If the birth terms *X*_*i*_ take the form *X*_*i*_ = *p*_*i*_ *H*({*A*_*k*_}, {*B*_*k*_}), where *p*_*i*_ is the population fraction of allele *i* and *H* is any function of the populations of either allele at any age (where the same function H appears in each *X*_*i*_), then only the two competitive exclusion steady-states are attainable and the quantity

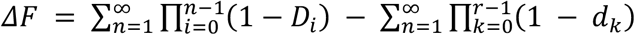

determines which one is stable, as stated in Eq (2) (see Appendix 1 in the Supplementary Information). This quantity corresponds to the difference in survival to age *n* summed over all ages *n* > 0. If Δ*F* is positive, allele *A* wins. If it is negative, *B* wins. Because it determines the result of the competition between the two alleles, Eq (2) acts as a fitness function for our competition.

Since Δ*F* is a fitness-like quantity, we can use it to judge the affinity of exponentially and linearly reproducing populations for longevity and vitality by studying the fitness of those traits on the competing alleles. To do so, we first specify reproduction functions that produce exponential and linear growth and verify that they are of the form *X*_*i*_ = *p*_*i*_ *H*({*A*_*k*_}, {*B*_*k*_}), so that Eq (2) applies. For linear growth, we assume a single queen produces a constant number *g* of offspring during each interval *dt*. For simplicity, we assume the allele is passed down purely through the father (removing the queen effects), an equal sex ratio, and random mating. Under these assumption, each phenotype contributes offspring in proportion to their current frequency in the population. Thus if A(t) and B(t) are the total count of allele A and B in the population, the population growth follows

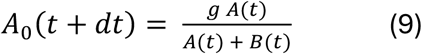

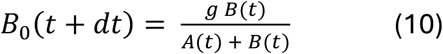

where *A*_0_ and *B*_0_ are the allele counts among the offspring (individuals of age 0). These growth terms satisfy the prerequisites for Equation (2).

For an exponentially growing population, the death probabilities are constant, so the birth term must saturate, or the population will either grow indefinitely or collapse. Given the aforementioned assumptions, the growth rate is

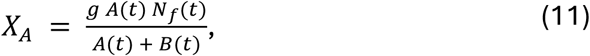

where *N*_*f*_(*t*) = *N*(*t*)/2 is the total number of females in the population, *g* is female fecundity (matching the queen’s in the linear population), the factor 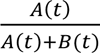 comes from random mating, and the factor *N_f_(t)* follows from the assumption of paternal inheritance. This growth rate is the same as the linear model, except that it is multiplied by the number of reproductive females. To keep the population finite, the factor *N(t)* must be replaced with a saturating function *S*(*N*(*t*)) = *S*(*A*(*t*) + *B*(*t*)). For reasons explained in Appendix 1 of the Supplementary Information, the form

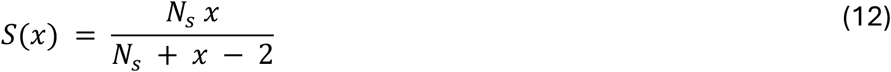

is an appropriate choice. Here, *N_s_* characterizes the population size at which reproduction saturates. Our full logistic growth model is then

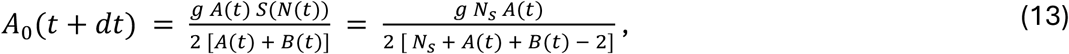

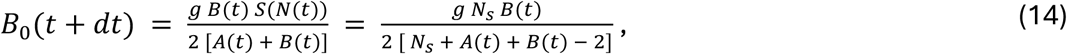

Once again, we can see that *X*_*i*_ = *p*_*i*_ *H*({*A*_*k*_}, {*B*_*k*_}), so Eq (2) to serves as the fitness function.

With Equations (2), (9-10), and (13-14), we can determine the outcome of competition between alleles differing in vitality and longevity for linear and exponential reproduction separately. However, our goal is to compare those outcomes across these growth modes. We have already formulated our growth laws so that the reproduction term of the linear and exponential populations match. We must also ensure that the two populations experience the same environment. To do this, we require that their total population sizes at steady state be equal, so that both have the same carrying capacity—that is, both are constrained by an environment that can sustain the same total number of individuals.

If the death probabilities are the same in the linear and exponential populations, then the exponential population will always be larger, because its birth rate equals the linear growth birth rate times the (effective) total number of females. We therefore adjust the age-independent component *γ* of the hazard rate to equalize the carrying capacities of the two populations without altering the age-dependent components. The adjusted value, *γ*’, is determined numerically so that the exponential growth model has the same steady-state population size as the linear growth model. Appendix 1 of the Supplementary Information gives expressions for these steady-state population sizes and shows that, under this matching, the exponentially growing population has a lower average age than the linearly reproducing one.

Finally, we can compare the affinities for the two alleles between the exponential and linear populations. Using SciPy’s *fsolve* root-finding algorithm, we numerically searched the vitality-longevity parameter space to find the values of (*x*, *y*) for which Δ*F* = 0. These roots correspond to points where both alleles have equal fitness. We did this procedure for a linear population and an equivalent exponential population (see details in Appendix 1 in the Supplementary Information), yielding the fitness nullclines in Figure 2. This analysis revealed that the linear population is more sensitive to changes in longevity than the exponential population.

### Colony reproduction

We extend the analytical model of the previous section to enable colony reproduction in order to test whether exponential growth of colonies caused a selective force for vitality in opposition to the selective force for longevity observed in the individual-reproduction models. To keep the analysis tractable, we focus on a specific case of colony reproduction described below, leaving more general cases to the numerical model (Results 2b). This case is particularly simple and requires no new derivations, only the application of results already obtained.

Here, we assume that allele fixation within a colony is generally fast compared to colony birth/death, so we can regard each colony as either a colony of allele A or allele B. (This assumption is supported by our numerical results, where most colonies were fully dominated by a single allele.) Each such colony seeds other colonies of the same genotype, and these colonies can collapse and "free up" a colony site. Colony seeding and collapse can be regarded as birth and death processes. In this way, we have now modeled colony reproduction exactly as an exponentially reproducing population, as in Equations (3-6) and (13-14).

Now, however, the death rates are the collapse rates of the colonies, not the death rates of the individuals. Thus, the parameters that would feel the selective force for vitality are the death probability parameters of the colonies, not the Gompertz parameters of the individuals. These are, in general, completely independent parameters; the colony death rate as a function of its age—i.e. time since founding—does not have to be related to the age-dependent death rates of its members. Thus, when properly applied, there is no competing force on the colony level selecting for vitality. Without other mechanisms explicitly linking these parameters, the linearly growing population’s preference for longevity is unaltered.

While exponential colony reproduction does not constitutively negate the eusocial population’s sensitivity to longevity, it is possible that colony reproduction could select against longevity through mechanisms unrelated to the growth-rate-dependent selective force studied here. We now use a simple model of allele competition among individuals in reproducing colonies to show that there is no such built-in selection against longevity.

Additional mechanisms—peculiarities in the way the species founds and disbands colonies, ones explicitly acting against longevity—would be required to negate this effect.

We now show that colony selection depends on the ratio of seeding rate to collapse rate. Let *r* be the number of available colony sites, *A* the number of colonies of type A, and *B* the number of colonies of type B. If there are *M* total colony sites on a landscape, and we assume colony collapse happens at a constant per-colony (allele dependent) rate, and the per-colony seeding rate is proportional to the number of available sites, then the competition for colony sites is captured by the equations

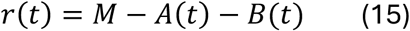

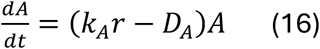

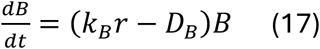

There are only 3 possible steady-states of this system, given by

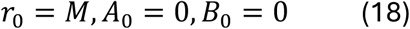

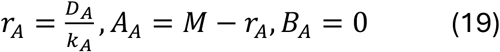

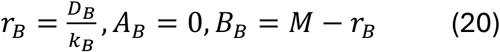

One is the trivial steady-state with no colonies, and the other two obey competitive exclusion. The only stable steady-state is the one with the lowest value of *r* (states with higher *r* values are unstable to invasion) [70].

Since the intra-colony dynamics are fast, the colony seeding and collapse rates are functions of the colony size and the steady-state age distribution across colony members. Consider the special case where these rates depend only on colony size. Bigger colonies should have a higher seeding rate (more individuals available to found a new colony) and a lower collapse rate (better ability to weather tough periods, lower probability of stochastic fluctuations to collapse). Thus, whichever allele supports larger colony size will be selected for, since it has a higher birth rate *k* and lower death rate *D* (of the colonies), and thus has the lower value of *r* at steady-state. In Appendix 1 of the Supplementary Information, Equation (S.23), we showed that the steady-state population of a linearly reproducing colony was an increasing function of our fitness parameter from Equation (1). Therefore, the allele that is selected for on the colony level is the exact same allele that is selected for on the individual level, with the exact same fitness function.

This shows that, at least under these simplifying assumptions, colony reproduction has no effect on selection for longevity in eusocial populations. Exponential reproduction of the colony alone cannot change the direction of selection. Such directional changes, if they occur, may require mechanisms linking the individual phenotypes to colony-level parameters, either through the colony’s own age-dependent dynamics, or through the dependence of the seeding and collapse rates on the underlying age distribution of the individuals.

### Numerical models

#### 1. The NMR model

##### A. Individual Reproduction

This is an agent-based model in Python [71], version 3.13.2. Individuals are diploid male or female and sex frequencies average 50%. In exponential populations, mating between males and females is random. In eusocial/linear populations, there is only one mating female, the queen. Upon death of the queen, she is replaced randomly by another female. Genes assort in the usual way by Mendelian genetics, and genes can be recessive or dominant. Time in the simulation proceeds in steps called “ticks”.

The model balances births against deaths. Births are governed by parameters including age at sexual maturity, probability of reproduction, and litter size. Deaths are governed by age-dependent mortality according to the Gompertz law (see Appendix 2) and by a logistic equation that increases mortality with increased population size. As populations grow, density-dependent mortality increases, and populations come to an equilibrium size with births balancing deaths. We introduce various alleles that affect parameters of the Gompertz equation and study their effects. These “Williams alleles” [34] influence the α and β parameters of the Gompertz equation in opposite directions. Typically, we compete one allele with high vitality (low α, high β) against another with high longevity (high α, low β) and observe which allele becomes fixed in a simulated population under linear or exponential reproduction. See Appendix 2 in the Supplementary Information for details.

##### B. Colony Reproduction

Colony reproduction uses the same underlying code as individual reproduction (part A above), but super-imposes the reproduction of colonies. There are two modes of colony reproduction, distinguished by different parameter settings.

###### i. Ant mode

In Ant mode, new colonies are established by single queens carrying sperm from a mating flight. These give birth and die by the rules of individual reproduction. Mating occurs in intermittent mating flights, not within colonies. When the queen dies, the whole colony dies, as in real ants. The probability of a new queen (formed during a mating flight) establishing a new colony is set by a logistic function that includes the existing number of colonies. The death of queens (and thus of colonies) is balanced by the formation of new colonies, eventually reaching an equilibrium population, which is at equilibrium both in terms of the number of individuals, and the number of colonies.

###### ii. Mole-rat mode

In Mole-rat mode, new colonies are established by groups of randomly chosen individuals from established colonies. One female becomes the queen and mates with males within the colony. The queen reproduces and dies by the same rules of the individual-reproduction model. When the queen dies, she is replaced (after a delay set by the “TICKS_TO_QUEEN_REPLACEMENT” parameter) by another female from the same colony. At intervals, “pioneers” leave to form new colonies. The probability of successfully forming a new colony is set by a logistic function that depends on the existing number of colonies. To allow colony turnover, colonies also face a small probability of catastrophic failure (typically 0.045). When a disaster occurs, there is a high probability of individuals in the colony dying and the colony may collapse. Loss of colonies through disasters is balanced by the formation of new colonies by pioneers.

Further details are provided in Appendix 2 and also in the “README” file posted with the code [72] where the NMR model is referred to as the “Health-Points-Model”. Further details can be requested from B.S. or B.F.

#### 2. The Queen effects-only model

To isolate the influence of the queen effects, we compare longevity and vitality alleles in exponential populations with similar size and age distribution. We simulate a population initially monomorphic for a resident allele. Once the age distribution reaches a steady state, we introduce a small number of individuals carrying a competing allele which confers lower longevity but higher vitality (i.e. higher *β* and lower *α*). We consider a range of invading alleles with the same reduced longevity but varying vitality. For each candidate allele, we repeatedly simulate its invasion and estimated the fixation probability as the proportion of simulations where the invader fixed. This observed fixation probability is then compared to the expected fixation probability of a neutral allele, equal to its initial frequency. This model was written in the R programming language [73].

We defined the fixation bias (FB) of the invading allele as FB = *f*/*p*_0_ − 1, where *f* is the estimated fixation probability and *p*_0_ is the initial frequency of the allele, calculated as the number of introduced mutant individuals divided by the total population size at the time of invasion. A positive FB indicates that the allele fixes more often than expected under neutrality (positive selection), while a negative FB indicates the opposite. Invading alleles with FB = 0 are therefore those whose vitality advantage exactly offsets their reduced longevity. Between the two population types, the one whose FB = 0 allele has the lower longevity is the population with stronger selection for longevity. This measure is robust to differences in effective population size between the two population types. Effective population size influences fixation probabilities—and therefore the magnitude of FB—but it does not change whether an allele fixes more or less often than expected under neutrality. Thus it does not affect which allele satisfies FB = 0.

The code for the Queen effects-only model is publicly available [74]. At this site it is referred to as the Gompertz-Mortality-Model.

## Author contributions

BF conceived of the study. BS and BF wrote and analyzed the NMR numerical model. RD wrote and analyzed the Queen effects-only computational model. CDK performed the analytical treatment. BF, RD, and CDK planned the manuscript. BF wrote the manuscript with contributions from RD and CDK. All authors contributed to manuscript edits and revisions.

## Acknowledgements

We thank Janet Leatherwood, Steve Skiena, Ken Dill, Liliana Davalos, Josh Rest, Krishna Veeramah, and Jeff Levinton for helpful discussions or advice on the manuscript. BF thanks Curtis Strobeck for assistance with aspects of population genetics. This work was funded in part by NIH R01s GM127542 and GM132238 (to BF).

