## Supplemental Information for "Eusocial reproduction selects for longevity"

### Appendix 1. Analytical model

Our analytical model wants to find the results of the competition between a wild-type allele and a Williams' allele to assess what Williams' allele has fitness zero. To do so, we need to model two populations. One is the collection $A_i(t)$ of wild-type individuals of age *i* at time *t*, the other is $B_i(t)$ of mutant individuals of age *i* at time *t*. To properly account for the age structure, it is easiest to use a discrete-time model with time step *dt* and ages measured in units of *dt*. Given the current populations at time *t*, the populations at the next time step *t*+*dt* are given by the rules

$$A_0(t + dt) \ = \ X_A \ , \tag{S.1}$$

$$A_{n+1}(t + dt) \ = \ (1 \ - \ D_n) \ A_n(t) \ , \tag{S.2}$$

$$B_0(t + dt) \ = \ X_B \ , \tag{S.3}$$

$$B_{n+1}(t + dt) \ = \ (1 \ - \ d_n) \ B_n(t) \ , \tag{S.4}$$

where $X_{A,B}$ is the amount of births of either type during the period *dt*, and individuals of age *n* survive to age *n*+1 (again, measured in *dt*) with probability (1 - $D_n$) for the wild-type and (1 - $d_n$) for the mutant. For simplicity, we assume that the death probabilities $D_n$ and $d_n$ are constants that do not depend on the populations.

This set of equations is very general, able to capture many scenarios. We now need to specify how to get linear and exponential growth out of it. To do so, we will make a few assumptions. First, linear growth. For linear growth, we assume there is one queen that produces a constant number g of offspring during the interval dt. Thus, our model tracks how an allele would go about fixing within a single eusocial colony. Additionally, for simplicity, we assume that the allele is passed down purely through the father (removing the queen effects that we discussed in the main text), that the M:F ratio is exactly equal, and that mating is random. These last assumptions allow us to say that of the *g* individuals born during each *dt*, the fraction of the total for each phenotype is exactly the current population fraction of that phenotype. Denoting the total number of wild-type as *A(t)* and the total number of mutants as *B(t)*, we find that the linear population can be modeled by the equations

$$A_0(t + dt) \ = \ \frac{g \ A(t)}{A(t) + B(t)} \, , \tag{S.5}$$

$$A_{n+1}(t + dt) \ = \ (1 \ - \ D_n) \ A_n(t) \ , \tag{S.6}$$

$$B_0(t + dt) \ = \ \frac{g \ B(t)}{A(t) + B(t)} \, , \tag{S.7}$$

$$B_{n+1}(t + dt) \ = \ (1 \ - \ d_n) \ B_n(t) \ . \tag{S.8}$$

For the exponential population, since the death probabilities are constant, we must use a birth term that saturates. Otherwise, the exponential population will either blow up to infinity or decay to zero. For pure exponential growth with random mating, equal M:F ratio, females reproducing at the same rate as our linear growth queen, and father-only genetics, the growth rate would be

$$X_A = \frac{g\,A(t)\,N_f(t)}{A(t)+B(t)}, \quad (S.9)$$

where $N_f(t) = N(t)/2$ is the total number of females in the population ($g$ comes from each female being the same as the queen, while the population fraction of *A(t)* comes from random mating, while the total female population $N_f(t)$ comes from father-only genetics). It is the same as the linear model's growth rate, except now it is multiplied by the number of females that can reproduce.

The factor of *N(t)* must be replaced with a saturating function *S(N(t)) = S(A(t) + B(t))* to keep the population finite. We are essentially saying that we use an effective population of size *S(N(t))* instead of the full population. This saturating function should satisfy three conditions: (1) *S(x)* is an increasing function of its argument (adding more individuals should not decrease the reproduction rate), (2) when there are only two individuals in the total population, *S*(2) = 2, so that a population with one female reproduces at rate *g*, the same as the queen in the linear model (which means that we are using an exponential growth model matched to our linear population), and (3) *S(x)* should be proportional to *x* when *x* is small (it reduces to exponential growth), but it should have a horizontal asymptote as *x* increases (a finite steady-state exists). The Michaelis-Menten function

$$S(x) = \frac{N_S\,x}{N_S + x - 2} \quad (S.10)$$

satisfies these conditions, where $N_s$ is an intrinsic phenomenological parameter (i.e. it can be measured experimentally) of the species that characterizes when the reproduction saturates. Further, it is easy to see that its Taylor expansion for small population values is first exponential growth (one term), then logistic growth (two terms). Our full logistic growth model is then

$$A_0(t+dt) = \frac{g\,A(t)\,S(N(t))}{2\,[A(t)+B(t)]} = \frac{g\,N_S\,A(t)}{2\,[\,N_S + A(t) + B(t) - 2]}, \quad (S.11)$$

$$A_{n+1}(t+dt) = (1 - D_n)\,A_n(t), \quad (S.12)$$

$$B_0(t+dt) = \frac{g\,B(t)\,S(N(t))}{2\,[A(t)+B(t)]} = \frac{g\,N_S\,B(t)}{2\,[\,N_S + A(t) + B(t) - 2]}, \quad (S.13)$$

$$B_{n+1}(t+dt) = (1 - d_n)\,B_n(t). \quad (S.14)$$

#### The Gompertz form of the death probabilities

We now need to find the form of the death probabilities that gives us the Gompertz law for the death rate. For the Gompertz-Makeham law, the per-capita death rate (hazard) of the population is

$$\mu(t) = \alpha\,e^{\beta t} + \gamma. \quad (S.15)$$

(Note: in the main text, we used h(t) for the hazard; we are now switching to $\mu(t)$ to underscore the fact that it is a death rate). Now assume we have a population *N(t)* of individuals that are age 0 at *t* = 0. We need to find $1 - D_n = N(n+dt)/N(n)$, the fraction of individuals alive at age *n* that are still alive at age *n* + *dt*. To do so, we note that

$$\frac{dN}{dt} = -\mu(t)\,N(t), \quad (S.16)$$

so that we can solve for *N(t)*:

$$N(t) \ = \ N(0)\, e^{-\int_0^t dt'\, \mu(t')} \ . \qquad \text{(S.17)}$$

Evaluating, we find

$$1 \ - \ D_n \ = \ \frac{N(n+dt)}{N(n)} \ = exp\,[-(\frac{\alpha}{\beta}) \ \ exp\,(\beta n)\,(exp\,(\beta\, dt) \ - \ 1) \ - \ \gamma\, dt]\ . \qquad \text{(S.18)}$$

We can verify that *N(n + dt)* ~ *N(n)* + *dt* (d*N(n)*/dt) matches the result of our discrete time model, *N(n + dt)* = (1 - $D_n$) *N(n)* when *dt* is small by showing that

$$\lim_{dt\,\to 0} D_n/dt \ = \ \mu(t) \ \ . \qquad \text{(S.19)}$$

So, we have found the form of the death probabilities that give us the Gompertz-Makeham mortality law. Each phenotype consists of a choice of $\alpha$, $\beta$, and $\gamma$, implying a fixed Gompertz curve.

#### Enforcing a common carrying capacity

Next, we need to ensure that our linear and exponential populations are seeing the same environment. To do so, we force their total populations at steady-state to be the same, so that they have the same carrying capacity. In the models we have written down, if the death probabilities are the same for the linear and exponential populations, then the exponential population will always be higher because its birth rate is the linear growth birth rate times the (effective) total number of females. Therefore, we must use a different value of $\gamma$ (the age-independent death rate) to make the carrying capacities the same without introducing any bias into the age-dependent parts of the death rate in the exponential population relative to the linear population. This new value of $\gamma$, call it $\gamma$', will have to be found numerically to ensure that the steady-state value of the total population for the exponential growth model is the same as the linear growth model.

We know that $\gamma$' > $\gamma$. What is the consequence? Consider for the moment just a single linear population, with no mutant. It is described by

$$A_0(t \ + \ dt) \ = \ g\ , \qquad \text{(S.20)}$$

$$A_{n+1}(t + dt) \ = \ (1 \ - \ D_n)\, A_n(t)\ , \qquad \text{(S.21)}$$

where for this section only *A* refers to a linear wild-type and *B* refers to an exponential wild-type, not a mutant. The steady-state age distribution can be found by setting $A_i(t+dt) = A_i(t)$. This condition implies that

$${A_0}^{SS} \ = \ g\ , \qquad \text{(S.22)}$$

$${A_n}^{SS} \ = \ (1 \ - \ D_{n-1})\, {A_{n-1}}^{SS} \ = \ g\, \prod_{i\,=\,0}^{n\,-\,1}(1 \ - \ D_i) \ \ . \qquad \text{(S.22)}$$

Summing these up, the total population at steady-state will be

$$A^{SS} \ = \ g\,[1 \ + \ \sum_{n=1}^{\infty} \prod_{i=0}^{n-1}(1 \ - \ D_i)]\ . \qquad \text{(S.23)}$$

and the average age is

$$\langle age \rangle \;=\; \sum_{n=0}^{\infty} \frac{n A_n{}^{SS}}{A^{SS}} \;=\; \frac{\sum_{n=1}^{\infty} n \prod_{j=0}^{n-1}(1-D_j)}{1+\sum_{m=1}^{\infty}\prod_{i=0}^{m-1}(1-D_i)} \;. \tag{S.24}$$

Since the factor of *g* cancels, the average age only depends on the death probabilities (which makes sense: the age distribution has one degree of freedom for each possible age; once you know $A_0$, the rest of the distribution is fixed by the death probabilities; the number of births sets the total population level, but does not impact the relative populations at each age).

For a single exponential population, the model is

$$B_0(t+dt) \;=\; \frac{g\, N_s\, B(t)}{2[N_s + B(t) - 2]}\,, \tag{S.25}$$

$$B_{n+1}(t+dt) \;=\; (1 \;-\; D_n{}') \, B_n(t)\;, \tag{S.26}$$

where $D_n$' uses $\gamma$' instead of $\gamma$. At steady-state, we find

$$B_0{}^{SS} \;=\; \frac{g\, N_s\, B^{SS}}{2[N_s + B^{SS} - 2]}\,, \tag{S.27}$$

$$B_n{}^{SS} \;=\; B_0{}^{SS}\, \prod_{i=0}^{n-1}(1 \;-\; D_i{}')\;. \tag{S.28}$$

To find $B^{SS}$, we must sum:

$$B^{SS} \;=\; \frac{g\, N_s\, B^{SS}}{2[N_s + B^{SS} - 2]}\,[1 \;+\; \sum_{n=1}^{\infty}\prod_{i=0}^{n-1}(1 \;-\; D_i{}')]\;. \tag{S.29}$$

This equation is a quadratic for $B^{SS}$, with one root being $B^{SS}$=0. Dividing by $B^{SS}$ removes this root, so that we can find

$$B^{SS} \;=\; 2 \;-\; N_s \;+\; (\tfrac{g\, N_s}{2})[1 \;+\; \sum_{n=1}^{\infty}\prod_{i=0}^{n-1}(1 \;- D_i{}')]\;. \tag{S.30}$$

This value looks like it can be negative; in the cases where it is, then only the $B^{SS}$ = 0 steady-state exists, indicating that the population is not able to reproduce fast enough to survive.

To calculate the average age, we can see that $B_n{}^{SS}$ and $B^{SS}$ are proportional to $B_0{}^{SS}$. Accordingly, that factor cancels, so that our answer for the average age takes the same form as for the linear case above (unsurprisingly, since the death probabilities fix the distribution and the growth strategy only fixes the total population level):

$$\langle age \rangle \;=\; \sum_{n=0}^{\infty} \frac{n\, B_n{}^{SS}}{B^{SS}} \;=\; \frac{\sum_{n=1}^{\infty} n \prod_{j=0}^{n-1}(1-D_j{}')}{1+\sum_{m=1}^{\infty}\prod_{i=0}^{m-1}(1-D_i{}')} \;. \tag{S.31}$$

We know that $D_n$' uses $\gamma$' instead of $\gamma$. How does that change the average age? First, we need to write the average age as a function of $\gamma$. To do so, first note that

$$1 \;-\; D_n \;=\; e^{-\gamma\, dt}\, h_n(\alpha, \beta)\;, \tag{S.32}$$

which allows us to write

$$\prod_{i=1}^{n-1}(1 - D_n) = e^{-\gamma\, dt\, n}\, f_n(\alpha,\beta) \,, \qquad (S.33)$$

where $h_n$ and $f_n$ are functions that do not matter for the current point, other than that they do not depend on $\gamma$. Accordingly, the average age can be written as

$$\langle age \rangle = \frac{\sum_{n=1}^{\infty} n\, e^{-\gamma\, dt\, n} f_n(\alpha,\beta)}{1 + \sum_{m=1}^{\infty} e^{-\gamma\, dt\, m}\, f_m(\alpha,\beta)} \,. \qquad (S.34)$$

Now we can take the derivative of the average age with respect to gamma:

$$\frac{d\ \langle age \rangle}{d\gamma} = \frac{[1 + \sum_{m=1}^{\infty} e^{-\gamma\, dt\, m} f_m(\alpha,\beta)]\, [\sum_{n=1}^{\infty} -dt\, n^2\, e^{-\gamma\, dt\, n}\, f_n(\alpha,\beta)]}{[1 + \sum_{m=1}^{\infty} e^{-\gamma\, dt\, m}\, f_m(\alpha,\beta)]^2} -$$

$$- \frac{[\sum_{n=1}^{\infty} n\, e^{-\gamma\, dt\, n}\, f_n(\alpha,\beta)][\sum_{m=1}^{\infty} -dt\, m\, e^{-\gamma\, dt\, m}\, f_m(\alpha,\beta)]}{[1 + \sum_{m=1}^{\infty} e^{-\gamma\, dt\, m}\, f_m(\alpha,\beta)]^2} \qquad (S.35)$$

$$= -dt\, \frac{[\sum_{n=1}^{\infty} n^2\, e^{-\gamma\, dt\, n}\, f_n(\alpha,\beta)\,]}{[1 + \sum_{m=1}^{\infty} e^{-\gamma\, dt\, m}\, f_m(\alpha,\beta)]} + dt\, \frac{[\sum_{n=1}^{\infty} n\, e^{-\gamma\, dt\, n}\, f_n(\alpha,\beta)][\sum_{m=1}^{\infty} m\, e^{-\gamma\, dt\, m}\, f_m(\alpha,\beta)]}{[1 + \sum_{m=1}^{\infty} e^{-\gamma\, dt\, m}\, f_m(\alpha,\beta)]^2} \qquad (S.36)$$

$$= -dt\ \langle age^2 \rangle + dt\ \langle age \rangle^{2} = -dt\, Var(age) < 0 \,. \qquad (S.37)$$

So, we have found that the average age is a monotonically decreasing function of gamma. Therefore, the average age of the equivalent exponential population, which uses $\gamma' > \gamma$, will necessarily be lower than the average age of the linear population, satisfying the conditions of our hypothesis.

#### Finding the Nullclines for Williams' Alleles in Competition with the Wild-Type

To make a Williams' allele, we need to simultaneously increase alpha and decrease beta. So, we have a fixed wild-type, with $\alpha_{WT}$, $\beta_{WT}$, and $\gamma_{WT}$. An antagonistic mutant will have $\alpha_M > \alpha_{WT}$, $\beta_M < \beta_{WT}$, $\gamma_M = \gamma_{WT}$ (or, we can flip the signs). Then, for every $\beta_M$, we can try to find the $\alpha_M$ that causes no selection to happen between the wild-type and the mutant (i.e. the fitness of the mutant and the wild-type are the same). We can then map out the whole change in fitness nullcline for the linear population. For the exponential population, we do the same procedure, except we use the $\gamma_{WT}'$ that matches the linear model's carrying capacity, and we will find different values of $\alpha_M$ that offset our changes in $\beta$, i.e. a different nullcline curve. Our hypothesis is that the exponential growth nullcline will be shifted from the linear nullcline because it will take a smaller increase in alpha for the exponential population to offset the decrease in $\beta$. That is, the linear population is much more sensitive to changes in $\beta$, relative to changes in $\alpha$, than the exponential population is.

We must find the condition that tells us that no selection is happening between the wild-type and the mutant. Thus, we must analyze the competition between *A* and *B* in both models. We can do them both at the same time by noticing that they both take the form

$$A_0(t + dt) = p(t)\, H(\{A_i(t)\}, \{B_i(t)\}) \,, \qquad (S.38)$$

$$A_{n+1}(t + dt) = (1 - D_n)\, A_n(t) \,, \qquad (S.39)$$

$$B_0(t + dt) = [1 - p(t)]\, H(\{A_i(t)\}, \{B_i(t)\}) \, , \tag{S.40}$$

$$B_{n+1}(t + dt) = (1 - d_n)\, B_n(t) \, , \tag{S.41}$$

where $p(t)$ is the population fraction of the wild-type and $H$ is an arbitrary function of all the $A_i$ and $B_i$. We want to find $p^{SS}$, the steady-state values of the population fraction. We know two values already: $p^{SS} = 1$ occurs when the wild-type fixes in the population and the mutant dies out, just giving us the wild-type's steady-state age distribution; $p^{SS} = 0$ occurs when the mutant fixes in the population and the wild-type dies out, just giving the mutant's steady-state age distribution (note that since this model is deterministic, if fixation occurs, the more fit competitor will always fix in the population). To find any possible others (steady-states where fixation does not occur, but instead coexistence happens), we must solve for the steady-state values. The model's equations tell us that the steady-states satisfy

$$A_0{}^{SS} = p^{SS}\, H(\{A_i{}^{SS}\}, \{B_i{}^{SS}\}) \, , \tag{S.42}$$

$$A_n{}^{SS} = A_0{}^{SS} \prod_{i=0}^{n-1}(1 - D_i) \, , \tag{S.43}$$

$$B_0{}^{SS} = [1 - p^{SS}]\, H(\{A_i{}^{SS}\}, \{B_i{}^{SS}\}) \, , \tag{S.44}$$

$$B_n{}^{SS} = B_0{}^{SS} \prod_{i=0}^{n-1}(1 - d_i). \tag{S.45}$$

We can find an equation for $p^{SS}$ by calculating it directly using its definition:

$$p^{SS} = \frac{A^{SS}}{A^{SS} + B^{SS}} = \frac{p^{SS}\, H(\{A_i{}^{SS}\},\{B_i{}^{SS}\})[1 + \sum_{n=1}^{\infty}\prod_{i=0}^{n-1}(1 - D_i)]}{p^{SS}\, H(\{A_i{}^{SS}\},\{B_i{}^{SS}\})[1 + \sum_{m=1}^{\infty}\prod_{j=0}^{m-1}(1 - D_j)] + (1 - p^{SS}) H(\{A_i{}^{SS}\},\{B_i{}^{SS}\})[1 + \sum_{r=1}^{\infty}\prod_{k=0}^{r-1}(1 - d_k)]}. \tag{S.46}$$

Importantly, the factor of $H$ cancels from all terms, so we just end up with a relationship between $p^{SS}$ and the death probabilities:

$$p^{SS} = \frac{p^{SS}\, [1 + \sum_{n=1}^{\infty}\prod_{i=0}^{n-1}(1 - D_i)]}{p^{SS}[1 + \sum_{m=1}^{\infty}\prod_{j=0}^{m-1}(1 - D_j)] + (1 - p^{SS})[1 + \sum_{r=1}^{\infty}\prod_{k=0}^{r-1}(1 - d_k)]} \, . \tag{S.47}$$

If $p^{SS} = 0$ or $p^{SS} = 1$, this equation is automatically satisfied. Therefore, we can ignore these two cases and search for any other behavior. If we assume $p^{SS} \neq 0$, cancel the $p^{SS}$ that appears on both sides, and rearrange, we find the condition:

$$p^{SS}\, [\sum_{n=1}^{\infty}\prod_{i=0}^{n-1}(1 - D_i) - \sum_{r=1}^{\infty}\prod_{k=0}^{r-1}(1 - d_k)] =$$

$$= [\sum_{n=1}^{\infty}\prod_{i=0}^{n-1}(1 - D_i) - \sum_{r=1}^{\infty}\prod_{k=0}^{r-1}(1 - d_k)] \, . \tag{S.48}$$

This equation can only be satisfied if $p^{SS} = 1$ (which we already have accounted for) or the quantity in brackets is zero. Said another way, if the quantity in brackets is zero, then $p^{SS}$ can take on any value. This case is exactly when there is no selective difference between the wild-type and the mutant: neither is preferred to fix in the population. Therefore, the quantity

$$\Delta F = \sum_{n=1}^{\infty}\prod_{i=0}^{n-1}(1 - D_i) - \sum_{r=1}^{\infty}\prod_{k=0}^{r-1}(1 - d_k) \tag{S.49}$$

is a proxy for the fitness difference between the mutant and wild-type. If we plot the nullclines of $\Delta F$, then we are plotting the fitness nullclines. We can numerically find these nullclines by solving $\Delta F((1-x)\alpha,(1-y)\,\beta,\gamma) = 0$ and solving for $y$ given $\alpha$, $\beta$, $\gamma$, and $x$ for the linear population, then find the same with the appropriate $\gamma$' for the exponential population. It is the result of this procedure that is plotted in Fig. 2 in the main text.

### Appendix 2. Numerical models

#### Modeling the Gompertz law, death, and environmental carrying capacity: the NMR Model

In this model, animals are born with a certain "vitality" which is inversely related to the alpha parameter in the Gompertz equation. In the text, we refer to this quality as "vitality", but in the code it is symbolized as "HP" ("Health Points"). That is, higher HPs equates to higher vitality, lower $\alpha$, and higher survival. HPs have a half-life, which is set by the $\beta$ parameter in the Gompertz equation. In the code, we use the parameter R (for "Rate") which sets the half-life of HPs. In the text, we refer to this parameter R as "longevity". R is inversely related to the $\beta$ parameter of the Gompertz equation. That is, HPs decline exponentially with age (i.e., with "ticks", the measure of time in the code), with their half-life set by R.

Declining HPs on their own do not directly cause death. However, individuals in our model are subjected to randomly-drawn "insults" every "tick". The insults are drawn from a specific distribution of insult sizes (see below) quantified in negative HPs. If the number of negative HPs in the insult exceeds or equals the individual's HPs, the individual dies. Otherwise, the individual recovers (over a time in ticks set by the "IRT" parameter (Insult Recovery Time), typically 2). Each insult is drawn from a distribution of insults of various sizes. Small insults are common; large insults are rare. Since HPs decline exponentially with age, older individuals are exponentially more likely to encounter a deadly insult (see below).

Importantly, we have shown that when the sizes of insults are distributed as 1/x^2, where x is randomly drawn from the uniform distribution on the interval (0,1), this model of HP and Insults reproduces the Gompertz law for hazard (see Fig. A2-1).

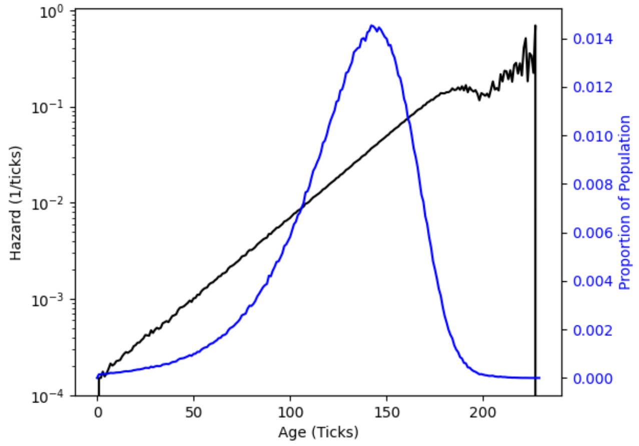

Fig. S-1. **In the NMR model, the increase in hazard is log-linear with time.**

The NMR model was run with typical parameters. Hazard (black line) is plotted as a function of age ("ticks"). Hazard is log-linear with time, consistent with the Gompertz equation. Noise at ages >200 is due to sampling, and has little or no effect on results because few or no individuals in populations have ages this high (blue line).

In summary, the model achieves a Gompertz mortality curve using an exponential decline in vitality (HPs) with age with a given half-life (set by R, longevity, the inverse of β), and imposing random insults whose magnitudes are drawn from a $\frac{1}{x^2}$ distribution, where $x$ is a properly parametrized random variable. We refer to this as a "Health Points and Insults" model.

Environmental carrying capacity is incorporated through a logistic function that scales insult size based on population size. As population increases, insults grow exponentially, modifying the randomly drawn insult magnitude to produce a final effective insult. At high population sizes, insults are large; at low population sizes, they remain small.

The logistic function is controlled by two parameters:

- pcE ("pop_cap_exponent") – sets the exponential rate at which insult size increases with population.
- pcM ("pop_cap_mult") – a scaling factor that adjusts insult size relative to individual Health Points.

These are scaled so that newborns can die from the largest random insults, but such events are rare.

An example of the Python code setting insult severity (using insult distribution T7, which is the one described above) is:

```
rand = random()*50000

insult_severity = 1000 / (rand**2)

population_size = len(CONFIG['alive_NMRS']['female']) +
len(CONFIG['alive_NMRS']['male']) if not after_pop_death else
CONFIG['InitialPopulation']

insult_severity = insult_severity * np.exp(CONFIG['pop_cap_mult'] *
(population_size**CONFIG['pop_cap_exponent']))
```

In this model, death is rare in small, young populations, which therefore grow in size. However, as populations grow, insult sizes and death increase because of the logistic equation, and at some population size, population size equilibrium is reached. The equilibrium population size is larger if the population is relatively young (because individuals have more Health Points), and the equilibrium size is smaller if the population is relatively old (because the individuals have fewer Health Points).

Table S1 shows parameters we commonly adjust in the model.

*Table S1. Key Parameters of the Health-Points model.*

| Parameter | Description | Common Values |
|---|---|---|
| **w** | Represents use of a "Williams" allele. A single genetic locus controls both vitality (HP) and longevity (R), with two possible alleles. | N/A |
| **HP** | Gene determining Health Points. Upper case ("HP") denotes dominant, lower case denotes recessive. Equivalent to vitality. | 900, 1000 |
| **R** | Gene determining Longevity. Sets the half-life of Health Points in ticks. Upper case ("R") denotes dominant, lower case ("r") denotes recessive. | 9, 10 (ticks) |

| | | |
|---|---|---|
| **pCE** | Population Cap Exponent. Controls how insult size increases exponentially with population size in the logistic equation. | 1.1 |
| **pcM** | Population Cap Multiplier. Adjusts the absolute size of insults. | 0.07 |
| **BP** | Birth Probability. Determines the likelihood of reproduction per tick. | 0.3 |
| **TTF** | Ticks Till Fertile. Number of ticks required to reach sexual maturity. | 3 |
| **IRT** | Insult Recovery Time. Number of ticks needed to recover from an insult. | 2 |
| **LS** | Litter Size. Maximum number of new agents born to a female per tick. | 1 |
| **Id** | Insult Distribution. Distribution of insult sizes. | T7 |
| **REPRODUCTION_TYPE** | Sets linear ("Lin") or exponential ("Exp") reproduction. | Lin, Exp |

Note that one locus has two phenotypes (HP and R), and the locus is either dominant (upper case) or recessive (lower case). Thus allowable genotypes are r10hp900 (our common recessive longevity allele), or R9HP1000 (our common dominant vitality allele). Other genotypes we have used include r9.2hp900 (the neutral longevity allele) and r9.3hp600 (the weak longevity allele). A genotype such as R10hp900 is nonsensical, since the entire allele has to be either dominant or recessive.

When Exponential and Linear populations are compared by simulation (e.g., Fig. 3), all parameters are the same for both populations. Only the mode of reproduction changes ("REPRODUCTION_TYPE=Lin" or "REPRODUCTION_TYPE=Exp") .

#### Colony Reproduction with the NMR model.

Colony reproduction uses the same underlying code as individual reproduction, but super-imposes the reproduction of colonies. There are two modes of colony reproduction, distinguished by different parameter settings.

Ant Mode. Each original colony (number of colonies is set by the "N0Colonies" parameter) starts with a heterozygous queen storing an infinite amount of sperm, 50% of each genotype. Ant queens do not mate within colonies. The ant queen gives birth and dies by the same rules as in individual reproduction. When the ant queen dies, the whole colony dies (as is the case in real ants). When the population of the colony reaches a threshold colony size (set by parameter "ThresholdColonySize") it sends out males and females ("reproductives") (but never the queen) who are excess to the threshold colony size. The males and females chosen to become reproductives could be chosen randomly from the colony, or they could be the youngest members of the colony, or they could be the oldest members of the colony (set by parameter "MatingFlightSelection"). These join other reproductives from other colonies in a mating flight, mate, and create new queens. New queens have a probability of founding new colonies; this probability is set by a logistic equation which includes the current number of colonies. As the number of colonies grows, it becomes exponentially less likely that a newly-formed queen is able to form colony. The population (both in number of individuals, and number of colonies) reaches equilibrium when queen deaths (which result in colony deaths) are balanced by formation of new colonies.

*Table S2. Key Parameters of Colony Reproduction*

| Parameter | Description | Common Values |
|---|---|---|
| **IMMORTAL_QUEEN_MODE** | Not used for these analyses. | False |
| **USE_COLONY_REPRODUCTION _MODE** | Toggles between individual reproduction and colony reproduction. | True, False |
| **N0Colonies** | Number of colonies in the landscape at the beginning of the simulation. | 16 |
| **MatingFlightInterval** | Number of ticks between "mating flights" per colony. | 6 |
| **ThresholdColonySize** | The population size a colony must reach before it sends out reproductives in a mating flight. The number of reproductives sent out is the number by which the current colony size exceeds this threshold. | 23 |
| **MatingFlightSelection** | How individuals are chosen for a mating flight. Usually set to "youngest", but "random" and "oldest" are other choices | youngest |
| **QueenHPMultiplier** | Not used for these analyses. Multiplies the Queen's HP, but not the HP of other individuals. | 1 |
| **BaseColonyFormationProbability** | Sets the base probability of formation of new colonies. This number is modified by the logistic cap on number of colonies. | 1 |
| **ColonyLogisticMult** | Helps set the logistic cap on number of colonies. | 0.06 |
| **ColonyLogisticExp** | The exponent in the colony logistic cap. | 1.1 |
| **PIONEER_GROUP_SIZE** | The number of pioneers needed to form a new colony. Must be set to "1" for Ant Mode | Ant Mode: 1<br>MR Mode: 8 |
| **COLONY_DISASTER_PROBABILITY** | Probability of encountering a "disaster". Must be set to "0" for Ant Mode. | Ant Mode: 0<br>MR: 0.045 |
| **COLONY_DISASTER_INSULT_MUL TIPLIER** | Multiplies severity of insults upon a disaster. Must be set to "1" for Ant Mode. Commonly 250000 for MR. | Ant Mode:1<br>MR: 250000 |
| **TICKS_TO_QUEEN_REPLACEMENT** | After death of a queen, number of ticks before a new queen in MR Mode. Must be "None" in Ant Mode. | Ant: None<br>MR: 20 |
| **MATES_WITHIN_COLONY** | Whether to allow mating within a colony. | Ant: False<br>MR: True |
| **REPRODUCTION_TYPE** | Linear ("Lin") or Exponential ("Exp") reproduction. Must be "Lin" in colony reproduction. | Lin |

**Python version and packages.** The results in the manuscript were obtained using Python 3.13.2, and the following packages: contourpy 1.3.3; cycler 0.12.1; fonttools 4.61.0; kiwisolver 1.4.9; matplotlib 3.10.7; numpy 2.3.5; packaging 25.0; pandas 2.3.3; pillow 12.0.0; pip 24.3.1; pyparsing 3.2.5; python-dateutil 2.9.0.post0; pytz 2025.2; scipy 1.16.3; six 1.17.0; tzdata 2025.2. With some other package versions, the logistic cap was altered—in one case, the number of colonies grew larger, and an unreasonable number of ticks were required for fixation.

### Isolating the Queen effect: the Gompertz Law Model

#### Exponential population dynamics

The population starts at carrying capacity with $K$ individuals at age 0, evenly split between males and females. Individuals are diploid, and all initially carry only the wildtype Williams allele. At each time step, all individuals undergo binomial mortality with probability of death depending on age and the size of the population as specified below. After the mortality stage, surviving females mate at random with surviving males and produce a Poisson-distributed number of offspring with mean fecundity $f_0$ offspring per female. Individuals are born at age 0 with an equal sex ratio, become reproductive at age 1, and inherit one allele from each parent. The mutant allele is dominant, so heterozygotes express the mutant phenotype. After a burn-in period to stabilize the age distribution, a proportion p0 of the population is randomly chosen to become homozygotes of the mutant Williams allele. After this invasion by the mutant allele, population dynamics continue until either allele fixes (i.e. excludes the other allele from the population entirely) or a set maximum number of years have passed since the start of the simulation.

#### Linear population dynamics

Here, the population is modeled as one single colony. The setup and dynamics are similar to that of the exponential population, except that all females except the Queen are sterile. Each time step, after the mortality stage, the Queen mates at random with one male and generates a Poisson-distributed litter with mean size equal to the carrying capacity $K$. If the Queen dies, she is immediately replaced at random by one of the other females in the population. After the burn-in period, the mutant allele is introduced to a random fraction $p_0$ of the population which may or may not include the Queen. As with the exponential population, after invasion by the mutant allele, population dynamics continue until either allele fixes or a set maximum number of years have passed since the start of the simulation. The parameterization used here ensures that average size and age distributions are similar between the Exponential and Linear populations.

#### Gompertz mortality

In both populations, mortality is density-dependent and increases with age following the Gompertz law. The hazard function—the instantaneous death rate at a given age conditional on survival up to that age—increases exponentially with age, $h(n) = \alpha\, e^{\beta n}$, where parameter $\alpha$ is the instantaneous death rate of newborns, while parameter $\beta$ relates to the aging rate, i.e., how quickly the hazard function increases with age. The probability that an individual of age $n$ in a population of size $N$ dies before reaching age $n+1$ is:

$$P_{\text{death}}(n, N) = 1 - \exp\left(-\frac{N}{K}\right) \exp\left(-\frac{\alpha}{\beta} \exp(\beta n)(\exp(\beta) - 1)\right)$$

Table S2 shows parameter values used in the simulations.

*Table S2. Parameter values used in the simulations.*

| Parameter | Value |
|---|---|
| Carrying capacity ($\boldsymbol{K}$) | 100 |

| | |
|---|---|
| Fecundity ($\boldsymbol{f_0}$) | 2 |
| Burn-in period [years] | 10 |
| Time step [years] | 0.2 |
| Maximum number of years | 1000 |
| Initial frequency of mutant allele ($\boldsymbol{p_0}$) | 0.05 to 0.5 by 0.05 increments |
| Number of replicates for exponential population | 1000 |
| Number of replicates for linear population | 5000 |
| $\boldsymbol{\alpha}$ parameter of the wildtype allele | 0.05 |
| $\boldsymbol{\beta}$ parameter of the wildtype allele | 0.15 |
| $\boldsymbol{\alpha}$ parameter of the mutant allele | 0.002 to 0.004 by 0.0005 increments |
| $\boldsymbol{\beta}$ parameter of the mutant allele | 1.2 |

### Appendix 3. Exact results for Fig. 5

Supplementary Table S1. Summary of Results from Numerical NMR Simulations

| Longevity Allele | Reproduction Mode | Longevity wins % (number) | Vitality wins % (number) | Neither |
|---|---|---|---|---|
| L10 V900 | Exponential | 6 (63) | 93 (928) | 9 |
| | Linear | 58 (584) | 42 (416) | 0 |
| | Colony (Mole-rat) | 96 (1893) | 4 (87) | 0 |
| | Colony (Ant) | 88 (875) | 12 (124) | 1 |
| L9.2 V900 | Exponential | 0 (2) | 99 (994) | 4 |
| | Linear | 52±3 (522) | 48 (478) | 0 |
| | Colony (Mole-rat) | 50±3 (595) | 49 (583) | 1 |
| | Colony (Ant) | 53.4±3.1 (534) | 46 (463) | 3 |
| L9.3 V600 | Exponential | 0 (0) | 100 (998) | 2 |
| | Linear | 48 (485) | 52 (515) | 0 |
| | Colony (Mole-rat) | 17 (341) | 83 (1659) | 0 |
| | Colony (Ant) | 42 (417) | 58 (580) | 3 |

These are the numbers used to generate Figure 5. The stated longevity alleles were competing with the standard L9 V1000 vitality allele. For the Linear, Mole-rat, and Ant versions of the L9.2 V900 simulations, we used Wald confidence intervals to find the error $\sigma$ of our measurement of the longevity win fraction:

$$\sigma = \frac{z}{\sqrt{n}}\sqrt{p(1-p)} \quad , \quad \text{(S.50)}$$

where n is the total number of runs, p is the fraction of longevity wins, and z = 1.96 for 95% confidence intervals.
